# Collective chemotactic localization emerges from interaction-driven phase transitions under temporally correlated noise

**DOI:** 10.64898/2026.05.10.724076

**Authors:** Gianna Arencibia, Martin Gutiérrez, Rafael Lahoz, Fivos Panetsos

## Abstract

Chemotactic microorganisms operate in environments where signals are not only noisy but temporally structured, with finite correlation times that can severely impair gradient sensing and target localization. While previous models have extensively characterized the effects of fluctuating environments on individual chemotaxis, most theoretical frameworks treat agents as non-interacting, leaving unresolved how inter-bacterial interactions reshape collective robustness under temporally correlated noise. Here, we introduce a two-dimensional agent-based model of interacting run-and-tumble bacteria navigating noisy chemotactic landscapes. We show that short-range isotropic cohesion induces a two-stage collective response: interactions first stabilize population connectivity and above a finite interaction threshold, this structural cohesion translates into robust target localization even in regimes where individual chemotaxis fails. The resulting transition reveals an intermediate phase of cohesive but weakly localized states, demonstrating that structural condensation and functional targeting are distinct collective observables. We further demonstrate that selective heterotypic interactions in binary populations produce a structurally distinct collective regime characterized by dynamically maintained red–blue contact networks composed of transient mixed dimers and local heterotypic motifs. Unlike isotropic cohesion, selective interactions reorganize local contact topology without generating macroscopic condensation. These structures are quantitatively characterized through bond density, mixing statistics, and anisotropy metrics, and are governed primarily by interaction specificity rather than by environmental noise persistence. Together, these results establish that collective chemotactic behavior is controlled by the interplay between temporal signal correlations and interaction topology. More broadly, in this work we identify collective localization and internal organization as partially independent emergent properties of interacting active matter under fluctuating environments.

## 1. Introduction

### 1.1 Chemotaxis in temporally structured environments

Chemotaxis, the directed movement of cells along chemical gradients, is one of the most fundamental and evolutionarily conserved mechanisms of biological motility. In bacteria, it enables navigation toward nutrients and away from harmful substances, supporting colonization, biofilm formation, and ecological competition^1,2^. In eukaryotic systems, analogous gradient-sensing processes underlie immune cell trafficking, wound healing, and morphogenesis, highlighting chemotaxis as a universal principle of directed biological motion^3,4^. Despite its ubiquity, a central unresolved question is how robust chemotactic navigation remains in environments where signals fluctuate over biologically relevant timescales. Natural chemical landscapes are inherently dynamic, shaped by diffusion, intermittent source activity, metabolic consumption, and transport across heterogeneous media^5,6^. Consequently, cells are exposed to signals that are not only noisy, but temporally correlated. These fluctuations introduce a characteristic persistence timescale, denoted here as the correlation time *τ*_*c*_, which quantifies how long environmental information remains coherent. Diffusion-limited transport, delayed secretion, and intracellular integration processes all contribute to generating signals with finite temporal memory. When fluctuations decorrelate rapidly, temporal sensing efficiently averages them out. However, when *τ*_*c*_ becomes comparable to or exceeds the intrinsic chemotactic response time of the cell, gradient estimation progressively deteriorates. In run-and-tumble systems, the relevant intrinsic timescale is the mean run duration *τ*_*run*_. The regime *τ*_*c*_ > *τ*_*run*_ is therefore expected to represent a fundamental limit of individual chemotactic sensing, because fluctuations persist across multiple sensing cycles and become indistinguishable from genuine gradients. In this regime, the limiting factor is no longer signal amplitude, but the mismatch between environmental memory and cellular processing. Understanding how collective interactions modify chemotactic behavior under these conditions constitutes the central motivation of this work.

### 1.2 Limitations of non-interacting chemotactic models

The quantitative description of bacterial chemotaxis is classically framed within the run-and-tumble paradigm, in which cells alternate between persistent runs and stochastic reorientation events, with tumbling rates modulated through temporal comparisons of sensed signals^8,9^. This framework successfully captures single-cell drift, macroscopic transport, and chemotactic efficiency in controlled gradients^2,10^. However, most existing chemotactic models implicitly assume that cells behave as independent, non-interacting agents. While this approximation is often sufficient in static or weakly fluctuating environments, it becomes fundamentally limiting in the presence of temporally correlated noise.

When environmental fluctuations persist over timescales longer than the characteristic run duration *τ*_*run*_, temporal sensing can no longer reliably discriminate genuine gradients from persistent stochastic perturbations. In models where fluctuations are represented as Ornstein–Uhlenbeck processes with correlation time *τ*_*c*_, chemotactic performance deteriorates sharply once *τ*_*c*_ > *τ*_*run*_: directional drift weakens, localization collapses, and populations disperse^7,11^. Importantly, this limitation is not merely quantitative but intrinsic to single-agent sensing. Because fluctuations remain coherent across multiple run-and-tumble cycles, increasing memory time alone cannot fully restore reliable navigation ^2,12^. In this regime, the breakdown of chemotaxis emerges as a collective-scale problem that cannot be resolved within purely non-interacting frameworks. This raises a central question: can inter-agent interactions generate collective mechanisms capable of stabilizing localization beyond the limits of individual chemotactic sensing?

### 1.3 Collective interactions as a route to functional robustness

A natural route to overcoming the limitations of single-cell chemotaxis lies in collective behavior. Active matter systems have shown that interacting populations of self-propelled agents can exhibit emergent order, enhanced transport, and robustness to fluctuations even when individual agents remain noisy or poorly directed^13,14^. Relatively simple interaction rules, including alignment or short-range attraction, are sufficient to generate coherent collective states and suppress stochastic dispersion at the population level^15,16^. Biological systems provide multiple examples of this principle. Collective chemotaxis, bacterial swarming and multicellular migration demonstrate that populations can outperform isolated cells in gradient sensing and directional persistence. Cooperative mechanisms such as density-dependent motility, quorum sensing and signal relay can amplify weak environmental cues and effectively average fluctuations across many interacting agents^17,18^. From this perspective, collective interactions may increase the effective signal-to-noise ratio available to the population as a whole^19^. Yet, whether cohesion alone can restore chemotactic function under temporally correlated noise remains unresolved. It is unclear whether inter-agent attraction simply stabilizes spatial organization or instead generates a genuinely collective localization mechanism capable of overcoming the breakdown of individual sensing. This distinction is critical because structural cohesion and functional targeting need not coincide: a population may form mechanically stable aggregates while remaining unable to localize near the signal source. Understanding how collective interactions transform chemotactic behavior in fluctuating environments therefore requires distinguishing between connectivity, condensation, and functional localization as separate emergent properties of the system.

### 1.4 From cohesion to interaction architecture

The simplest interaction capable of generating collective stabilization is isotropic short-range cohesion, in which pairwise forces depend only on inter-agent distance. Such interactions may arise from non-specific adhesion, surface peptides, or confinement within extracellular matrices, and provide a minimal physical mechanism for suppressing noise-driven dispersal. ^20,21^ However, cohesion alone does not guarantee function. A central premise of this work is that structural connectivity and chemotactic localization constitute distinct collective properties. Interactions may therefore stabilize mechanically cohesive groups without necessarily enabling efficient targeting of the signal source. This distinction suggests the existence of multiple collective regimes, including dispersed states, cohesive but weakly localized states, and fully localized collective phases. Biological interactions, moreover, are rarely isotropic. In many living systems, adhesion depends on cell identity, receptor ligand specificity, or differential affinity, generating selective interaction networks that actively reshape spatial organization^22,23^. The Differential Adhesion Hypothesis predicts that such selectivity can drive structured multicellular organization, including segregation, intermixing, and dynamically maintained heterogeneous assemblies^24^. In active systems, this raises a broader question: can interaction topology control collective architecture independently of chemotactic function? Under temporally fluctuating environments, selective interactions may reorganize local contact structure without necessarily enhancing localization itself. Interaction architecture and environmental memory therefore become coupled control parameters that jointly determine whether populations remain dispersed, condense into cohesive aggregates, or form dynamically maintained mixed-contact networks.

### 1.5 Scope and contributions of this work

In this work, we present a two-dimensional agent-based model of run-and-tumble bacteria navigating temporally correlated chemotactic landscapes. The framework extends the non-interacting LUCA1 model^25^ by introducing short-range inter-bacterial interactions while preserving the same temporal sensing dynamics. Environmental fluctuations are modeled as Ornstein–Uhlenbeck noise with tunable correlation time *τ*_*c*_, enabling systematic exploration of chemotactic behavior in persistence-dominated noisy environments. Two interaction architectures are investigated. In the isotropic model, bacteria interact through short-range peptide-mediated adhesion represented by a Morse-type potential. In the selective model, a binary red–blue population experiences heterotypic attraction and homotypic repulsion, allowing direct investigation of how interaction topology reshapes collective organization. We show that isotropic cohesion induces a two-stage collective response. Interactions first stabilize structural connectivity within the population and above a finite interaction threshold, this cohesion translates into robust collective localization even in regimes where individual chemotaxis fails. Importantly, structural condensation and functional targeting emerge as distinct collective observables, revealing an intermediate regime of cohesive but weakly localized states. We further demonstrate that selective interactions generate a qualitatively different collective regime dominated by transient heterotypic contact networks rather than macroscopic condensation. In this case, interaction specificity reorganizes local contact topology while leaving global localization only weakly affected, demonstrating a decoupling between internal organization and collective chemotactic function. Together, these results establish interaction topology and environmental memory as coupled control parameters governing collective chemotactic behavior under fluctuating conditions. More broadly, this work connects chemotaxis under temporally structured noise with non-equilibrium active matter, distributed biological self-organization, and programmable collective assembly. In particular, the emergence of robust collective states from local interaction rules parallels recent efforts in synthetic multicellular systems to engineer spatial organization through distributed signaling and irreversible local commitment, including programmable Voronoi-like microbial partitioning architectures^26^. This framework therefore provides a minimal physical basis for understanding how collective interactions reshape robustness, localization, and internal architecture in living active systems.

## 2. Model

To investigate how inter-bacterial interactions reshape chemotactic collective behavior under temporally correlated noise, we formulate five central hypotheses linking collective function, structural organization, and environmental fluctuations. The central premise of this work is that collective chemotaxis emerges not solely from individual sensing, but from the interplay between temporal noise structure and interaction architecture. H1. Interaction-driven rescue of collective localization. Moderate isotropic cohesion can restore collective localization in regimes where individual chemotaxis fails. We hypothesize that increasing adhesion strength *ε*suppresses noise-driven dispersal and stabilizes target occupancy collectively. However, this rescue is expected to be non-monotonic: weak interactions leave agents effectively independent, whereas excessively strong cohesion may stabilize mechanically coherent but poorly localized aggregates.

H2. Collective function is controlled by noise correlation time. The effectiveness of cohesion depends critically on the temporal persistence of environmental fluctuations. We therefore hypothesize that the correlation time *τ*_*c*_acts as a control parameter governing transition between dispersed, localized, and cohesive but weakly localized collective states. High structural connectivity is not expected to universally imply high chemotactic performance.

H3. Structural stabilization precedes functional localization. Collective rescue occurs through a sequential stabilization process. We expect isotropic interactions to first generate connected cohesive structures (*C*_*max*_ ↑) and only subsequently stabilize localization near the target (*η*_*ss*_ ↑). This predicts a decoupling between structural cohesion and functional chemotactic targeting.

H4. Selective interactions reorganize local contact topology. Selective heterotypic interactions reshape local contact organizations without necessarily inducing macroscopic condensation. In binary red–blue populations, heterotypic attraction combined with homotypic repulsion is expected to generate transient mixed-contact motifs characterized by enhanced heterotypic bond density and deviations from random mixing.

H5. Interaction architecture defines collective structural regimes. Isotropic and selective interactions generate qualitatively distinct collective architectures under identical chemotactic and noise conditions. Isotropic cohesion favors condensed cohesive phases, whereas selective interactions produce distributed heterotypic contact networks and frustrated mixed-contact states. Collective structure and collective function therefore emerge as partially independent properties of the system. Together, these hypotheses define a framework in which collective chemotaxis under fluctuating environments is governed by two coupled control principles: environmental memory and interaction topology. Within this framework, cohesion, localization, and internal organization emerge as distinct non-equilibrium collective phenomena with separate stabilization thresholds.

### 2.1 Overview

The aim of the model is to determine how inter-bacterial interactions modify collective chemotactic behavior under temporally correlated environmental noise, and specifically whether interaction-driven connectivity can restore localization in regimes where individual chemotaxis fails. The framework extends LUCA1 by introducing pairwise interactions while preserving the same run-and-tumble sensing dynamics. Two interaction architectures are considered. In the isotropic model, bacteria interact through short-range peptide-mediated adhesion, providing a minimal mechanism for collective cohesion. In the selective model, the population is divided into two subtypes with heterotypic attraction and homotypic repulsion, allowing investigation of how interaction topology reshapes collective organization. The model intentionally excludes direct gradient forces (*κ* = 0) and alignment interactions. Consequently, any emergence of localization, clustering, or internal structure arises exclusively from inter-agent coupling rather than from explicit collective steering mechanisms. This design enables a direct comparison between functional localization and structural organization under identical noisy chemotactic conditions.

### 2.2 Domain and agent dynamics

Simulations are performed in a two-dimensional square domain of side length *L* = 600 *μm*with reflective boundary conditions. Bacteria are modeled as self-propelled run-and-tumble agents characterized by position r_*i*_(*t*)and orientation *θ*_*i*_(*t*). The propulsion direction is

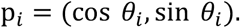

The dynamics of each bacterium follow

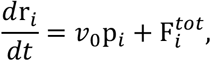

where *v*_0_ = 20 *μm*/*s*is the constant swimming speed and 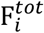 is the total interaction force exerted by neighboring agents. Time integration is performed using explicit Euler updates with timestep *dt* = 0.05 *s*, sufficiently small compared with all relevant dynamical timescales in the system.

Unless otherwise stated, simulations are performed with *N* = 200bacteria over a total duration *T* = 300 *s*, using *n*_*rep*_ = 8independent realizations per condition.

### 2.3 Signal landscape and temporal noise

The chemoattractant field is modeled as a static Gaussian source centered at the middle of the domain,

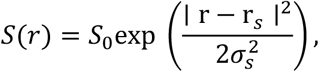

where *S*_0_ = 800is the signal amplitude, *σ*_*s*_ = 100 *μm*is the spatial width of the source, and r_*s*_ = (300,300) *μm*. To model fluctuating environments, each bacterium experiences an independent realization of temporally correlated Ornstein–Uhlenbeck (OU) noise. The perceived signal is defined as

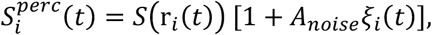

where *A*_*noise*_ = 0.35controls the fluctuation amplitude and *ξ*_*i*_(*t*)is an OU process with correlation time *τ*_*c*_. The present study focuses on the regime *τ*_*c*_ > *τ*_*run*_, where fluctuations remain correlated across multiple run-and-tumble cycles and temporal sensing becomes unreliable at the single-cell level. This persistence-dominated regime constitutes the baseline failure condition of LUCA1 and provides the central setting for testing whether collective interactions can restore localization.

### 2.4 Temporal sensing and run-and-tumble dynamics

Chemotactic navigation is implemented through temporal sensing. Each bacterium compares the current perceived signal to a stored memory value separated by a sensing interval Δ*t*_*s*_ = 0.5 *s*,

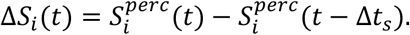

The tumbling rate is modulated according to

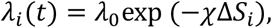

where *λ*_0_ = 1 *s*^−1^is the basal tumbling rate and χ = 0.005is the chemotactic gain. Positive temporal gradients reduce tumbling and promote persistent runs, whereas negative gradients increase reorientation probability.

At each sensing event, bacteria tumble probabilistically and adopt a new random orientation drawn uniformly from [0,2*π*). Between sensing events, orientations remain fixed except for reflective boundary collisions. This implementation preserves the temporal comparison logic of LUCA1 while explicitly incorporating fluctuating signal perception.

### 2.5 Isotropic peptide-mediated cohesion

To model cohesion mediated by surface-expressed peptides, we introduce short-range isotropic interactions using a Morse-type potential. The resulting interaction combines short-range repulsion with intermediate-range attraction, preventing overlap while allowing stable adhesive clustering. The interaction force depends exclusively on inter-particle distance and contains no angular or orientational dependence. Cohesion therefore emerges purely from isotropic attraction rather than from alignment or dipolar interactions. The characteristic interaction distance is *r*_0_ = 23 *μm*, with interaction range controlled by *a* = 0.08 *μm*^−1^. Forces are truncated beyond *r*_*cut*_ = 60 *μm* and clipped to a maximum magnitude *F*_*max*_ = 60 *μm*/*s* for numerical stability. The principal control parameter is the adhesion strength *ε*, explored over the range

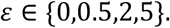

The case *ε* = 0corresponds exactly to the non-interacting LUCA1 limit.

### 2.6 Binary selective interaction model

To investigate how interaction topology reshapes collective organization, we extend the model to a binary population composed of red and blue bacteria. Each agent carries an identity label

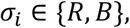

with equal numbers of red and blue particles unless otherwise stated.

Interactions depend on particle identity. Heterotypic pairs (R–B) experience attractive Morse-type interactions, whereas homotypic pairs (R–R and B–B) interact through purely repulsive short-range forces. This defines the interaction hierarchy: *RB*: attractive, *RR*: repulsive, *BB*: repulsive. The selective model represents effective complementary peptide or receptor-mediated binding between different bacterial subtypes. Unlike the isotropic system, where all contacts reinforce global condensation, selective interactions frustrate unrestricted aggregation and instead promote distributed mixed-contact structures. The relevant control parameters are the heterotypic adhesion strength *ε*_*RB*_, the interaction geometry (*r*_0_, *a*), and the homotypic repulsion scale.

### 2.7 Observables and order parameters

To characterize collective behavior, observables are grouped into three categories: localization, cohesion, and internal organization. This separation allows structural and functional aspects of collective behavior to be quantified independently and enables identification of regimes in which they become coupled, decoupled, or sequentially stabilized. Localization is quantified through the fraction of bacteria within a target radius around the signal source, the steady-state localization efficiency, and the mean tracking error. Cohesion is characterized by using cluster statistics, including largest-cluster fraction, number of fragments, and spatial dispersion. In the selective model, additional observables quantify internal organization, including heterotypic bond density, mixing index, contact topology, and cluster anisotropy.

To characterize collective behavior, observables are grouped into three categories: localization, cohesion, and internal organization. This separation allows structural and functional aspects of collective dynamics to be quantified independently and enables identification of regimes in which they become coupled, decoupled, or sequentially stabilized.

Localization observables quantify the ability of the population to accumulate and remain near the signal source. Cohesion observables characterize the emergence of connected collective structures independently of targeting performance. In the selective model, additional observables describe the topology and morphology of mixed-contact organization.

#### 2.7.1 Localization metrics

The instantaneous localization efficiency is defined as the fraction of bacteria contained within a target region of radius *R* = 100 *μm*centered on the signal source,

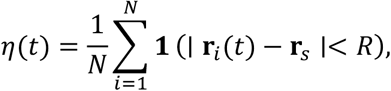

where **1**denotes the indicator function. Steady-state localization is quantified as the temporal average of *η*(*t*)over the final 20% of the simulation,

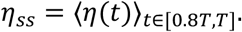

To distinguish localization from non-localized collective states, we define a localization threshold *η*_*th*_ = 0.20. The mean tracking error is additionally computed as

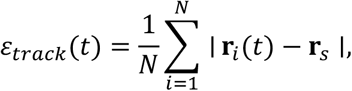

providing a continuous measure of the average population distance from the target.

Together, these observables quantify the functional chemotactic performance of the population under fluctuating environments.

#### 2.7.2 Cohesion and clustering metrics

Collective cohesion is quantified using distance-based clustering statistics. Two bacteria are considered connected whenever their separation is smaller than a threshold distance *r*_*c*_. Connected components are then identified to determine the instantaneous cluster structure of the population. From these connected components, we compute the size of the largest cluster *C*_*max*_(*t*), the total number of clusters *N*_*cl*_(*t*), and the fraction of agents belonging to the largest connected component,

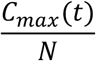

Spatial dispersion is quantified through the second central moment of particle positions,

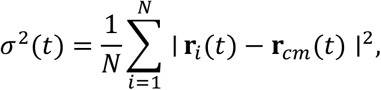

where **r**_*cm*_(*t*)is the instantaneous center of mass of the population.

These observables characterize structural connectivity independently of localization and allow identification of transitions between dispersed, cohesive, and condensed collective states.

#### 2.7.3 Selective structure metrics

For the binary red–blue system, additional observables quantify internal organization and contact topology. Contact is defined whenever two agents are separated by less than the interaction distance *r*_*c*_. We compute the number of heterotypic contacts *N*_*RB*_ and the numbers of homotypic contacts *N*_*RR*_ and *N*_*BB*_. The degree of intermixing is quantified through the mixing index

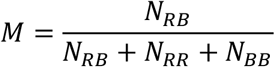

Values *M* ≈ 0.5 correspond to random mixing, whereas larger values indicate preferential heterotypic organization. To characterize cluster morphology, we compute the anisotropy parameter

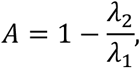

where *λ*_1_ ≥ *λ*_2_ are the eigenvalues of the covariance matrix of positions within the principal cluster. Compact isotropic clusters yield *A* ≈ 0, while elongated or chain-like structures produce *A* → 1. To further quantify selective organization, we additionally measure the mean chain length and the connectivity of heterotypic contact networks. These observables distinguish compact mixed condensates from distributed contact motifs, transient chains, and frustrated mixed states.

#### 2.7.4 Phase classification

Collective regimes are classified using the observables defined above. For isotropic interactions, three principal dynamical states are identified: dispersed states with weak connectivity, cohesive but non-localized states characterized by structural clustering without efficient targeting, and localized collective states in which cohesion and chemotactic accumulation coexist. For selective interactions, classification additionally incorporates mixing and morphology. Depending on cluster size, contact topology, and anisotropy, the system may exhibit dispersed or frustrated states, compact mixed-contact structures, or elongated heterotypic network-like assemblies.

### 2.8 Parameter summary

Unless otherwise stated, simulations use the following baseline parameters:

| Parameter | Symbol | Value | Description |
| --- | --- | --- | --- |
| Domain side | $L$ | $600 \mu\text{m}$ | Square domain with reflective boundaries |
| Number of agents | $N$ | 200 | Population size |
| Simulation time | $T$ | 300 s | Total simulation duration |
| Time step | $dt$ | 0.05 s | Euler integration timestep |
| Replicates | $n_{\text{rep}}$ | 8 | Independent realizations |
| Swimming speed | $v_0$ | $20 \mu\text{m/s}$ | Self-propulsion speed |
| Basal tumbling rate | $\lambda_0$ | $1 \text{ s}^{-1}$ | Mean run time $\tau_{\text{run}} = 1 \text{ s}$ |
| Sensing window | $\Delta t_s$ | 0.5 s | Temporal comparison interval |
| Chemotactic gain | $\chi$ | $0.005(\text{a.u.})^{-1}$ | Sensitivity to signal changes |
| Signal amplitude | $S_0$ | 800 | Peak signal intensity |
| Signal width | $\sigma_s$ | $100 \mu\text{m}$ | Gaussian source width |
| Target radius | $R$ | $100 \mu\text{m}$ | Localization region |
| Localization threshold | $\eta_{\text{th}}$ | 0.20 | Threshold for collective localization |
| OU correlation time | $\tau_c$ | 5 s | Temporal persistence of fluctuations |
| Noise amplitude | $A_{\text{noise}}$ | 0.35 | Relative signal fluctuation amplitude |
| Morse equilibrium distance | $r_0$ | $23 \mu\text{m}$ | Preferred adhesive spacing |
| Morse width | $a$ | $0.08 \mu\text{m}^{-1}$ | Interaction decay scale |
| Interaction cutoff | $r_{\text{cut}}$ | $60 \mu\text{m}$ | Maximum interaction distance |
| Force clamp | $F_{\text{max}}$ | $60 \mu\text{m/s}$ | Numerical stability bound |
| Adhesion strength | $\varepsilon$ | $\{0, 0.5, 2, 5\}$ | Isotropic cohesion strength |
| Signal-force coupling | $\kappa$ | 0 | No direct gradient force |

For the binary model, the additional interaction parameters are *ε*_*RB*_for heterotypic attraction and the repulsive scale for *RR*and *BB*pairs.

### 2.9 Numerical implementation

Pairwise interaction forces were evaluated at every timestep using vectorized distance calculations with a finite interaction cutoff. Although the interaction scheme scales nominally as *O*(*N*^2^), the effective computational cost remained modest because interactions were restricted to short ranges (*r*_*cut*_ = 60 *μm*) in a dilute 600 × 600 *μm* domain. Reflective boundary conditions were applied after each Euler update. Temporal signal memory was stored independently for each bacterium and updated exclusively at sensing events, ensuring faithful implementation of the finite temporal comparison window Δ*t*_*s*_. Unless otherwise stated, all reported quantities correspond to averages over *n*_*rep*_ = 8independent realizations and are presented as mean ±standard error of the mean (SEM).

### 2.10 Expected phenomenology

Based on the hypotheses formulated above, we expect collective behavior to emerge from the interplay between interaction strength and the temporal structure of environmental fluctuations. In the non-interacting limit (*ε* = 0), and particularly in the regime *τ*_*c*_ > *τ*_*run*_, temporally correlated noise is expected to impair temporal gradient discrimination, leading to weak or unstable localization. For isotropic interactions, we hypothesize that increasing adhesion strength produces a sequential collective transition. At intermediate interaction strengths, bacteria form cohesive connected structures that suppress noise-driven dispersal and stabilize localization near the signal source. However, structural cohesion and functional targeting are not expected to emerge simultaneously. Instead, we anticipate an intermediate regime characterized by cohesive but weakly localized states, indicating a decoupling between connectivity and chemotactic function. For selective interactions, we expect interaction topology to reorganize local contact structure without necessarily inducing macroscopic condensation. Heterotypic attraction combined with homotypic repulsion is predicted to generate dynamically maintained mixed-contact motifs and distributed heterotypic networks whose organization depends primarily on interaction specificity rather than on global localization efficiency. Consequently, isotropic and selective interactions are expected to produce qualitatively distinct collective architecture even under identical chemotactic and noise conditions. Together, these expected regimes provide a direct test of whether collective interactions can both stabilize chemotactic function and independently reshape internal organization under temporally structured noise.

## 3. Results

### 3.1 Breakdown of chemotaxis under temporally correlated noise

We first establish the baseline behavior of the non-interacting LUCA1 limit (*ε* = 0) to identify the conditions under which individual chemotactic sensing fails to sustain stable localization. In the absence of inter-agent interactions, collective organization emerges exclusively from temporal gradient sensing through run-and-tumble dynamics, providing the reference state against which the collective stabilization mechanisms introduced in this work can be evaluated. Under static or rapidly fluctuating signal conditions, populations robustly localize near the chemoattractant maximum. However, this robustness collapses when environmental fluctuations become temporally correlated over timescales comparable to or larger than the intrinsic run duration *τ*_*run*_. In this regime, temporal comparisons can no longer reliably distinguish genuine gradients from persistent stochastic perturbations.

For the baseline parameters used throughout this work (*S*_0_ = 800, *τ*_*c*_ = 5 *s, A*_*noise*_ = 0.35), the system operates in this persistence-dominated regime (*τ*_*c*_ > *τ*_*run*_). The resulting steady-state localization is *η*_*ss*_ = 0.189 ± 0.009(mean ± SEM, *n* = 6), below the localization threshold *η*_*th*_ = 0.20. The mean tracking error reaches *ε*_*track*_ = 200.9 ± 2.0 *μm*, close to the expected value for an approximately dispersed population in a 600 × 600 *μm*domain. Consistently, the largest cluster fraction remains low (*C*_*max*_ = 0.234 ± 0.016), reflecting only transient spatial proximity in the absence of cohesive interactions.

Comparison between different fluctuation spectra further demonstrates that chemotactic robustness depends primarily on temporal structure rather than on noise amplitude alone. White noise is efficiently averaged out by temporal sensing, whereas slowly varying fluctuations progressively bias gradient estimation. Ornstein– Uhlenbeck noise produces the strongest disruption because low-frequency fluctuations persist across multiple run-and-tumble cycles, effectively decorrelating the population from the target. Together, these results define the baseline failure regime of LUCA1. Once *τ*_*c*_ ≳ *τ*_*run*_, individual temporal sensing becomes insufficient to maintain robust localization despite the presence of a persistent mean gradient. This non-interacting failure states the central motivation for this work: whether collective interactions can rescue chemotactic localization beyond the limits of single-agent sensing.

### 3.2 Isotropic peptide-mediated cohesion rescues chemotactic localization under temporally correlated noise

We next investigated whether short-range isotropic cohesion can restore collective localization in the failure regime identified for the non-interacting LUCA1 model. Simulations were performed under persistent Ornstein– Uhlenbeck fluctuations (*τ*_*c*_ = 5 *s, A*_*noise*_ = 0.35), where individual chemotaxis alone is insufficient to maintain stable target localization. Introducing isotropic adhesion qualitatively reshaped the collective dynamics (Fig. 1). In the absence of interactions (*ε* = 0), the population remained only weakly localized, fluctuating near the localization threshold. Weak cohesion (*ε* = 0.5) produced only modest improvement. However, at intermediate interaction strength (*ε* = 2), the system underwent a pronounced collective rescue transition: the localization fraction increased rapidly and stabilized at *η*_*ss*_ ≈ 0.61, substantially above the non-interacting baseline. This enhancement was accompanied by a marked reduction in tracking error and the emergence of compact cohesive structures localized near the signal source (Fig. 1C–D). Importantly, the effect was strongly non-monotonic. Increasing adhesion further (*ε* = 5) did not improve localization and instead reduced long-term target occupancy despite maintaining cohesive aggregates. Representative snapshots show that strongly adhesive populations frequently formed mechanically stable clusters displaced from the source. These results demonstrate that cohesion and localization are distinct collective observables. Although increasing adhesion generally promotes structural cohesion, maximal connectivity does not correspond to maximal chemotactic performance. Instead, localization is optimized within an intermediate interaction regime where cohesion suppresses noise-driven dispersion while preserving sufficient collective mobility to remain responsive to the chemotactic landscape. Together, these findings establish that isotropic peptide-mediated interactions can rescue collective chemotaxis under temporally correlated noise, but only within a finite interaction window. Cohesion therefore acts not simply as a clustering mechanism, but as a collective stabilization process that transforms unreliable individual sensing into robust population-level localization.

**Figure 1.**
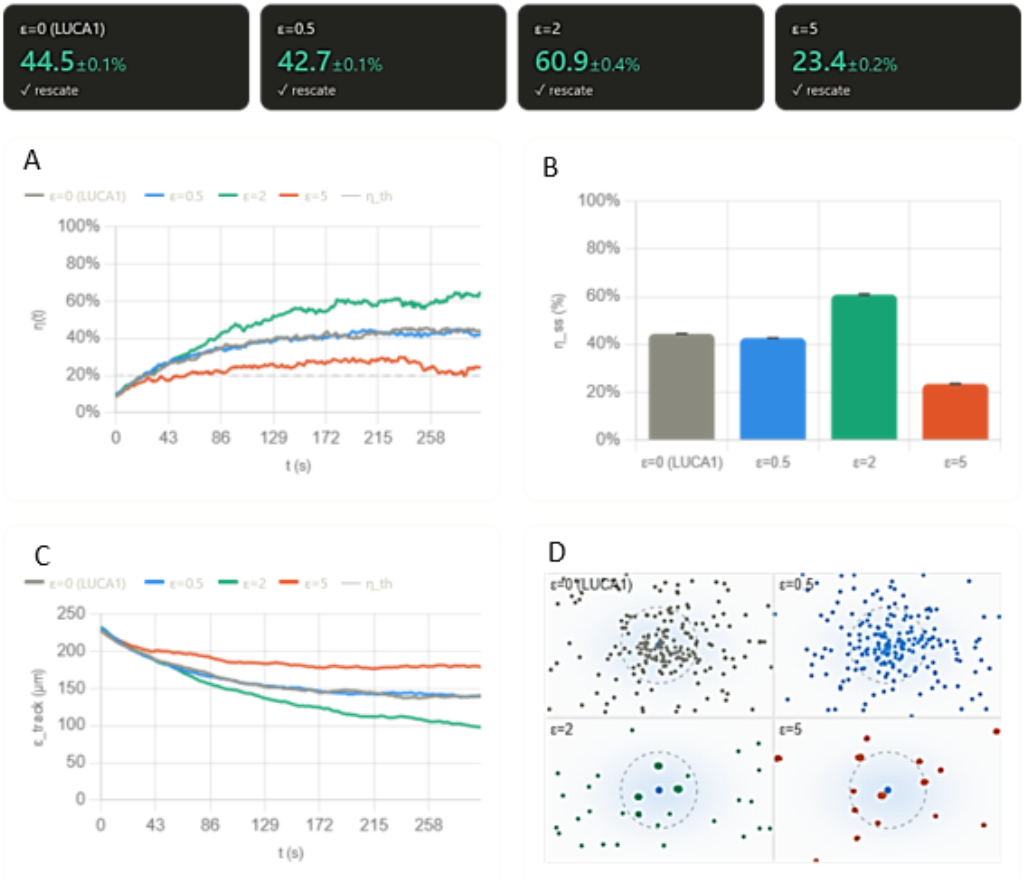
Isotropic cohesion rescues collective localization under temporally correlated noise. (A) Time evolution of localization efficiency η(t)for increasing adhesion strength ε. Dashed line: localization threshold η_th_ = 0.20. Intermediate adhesion (ε = 2) produces robust collective localization, whereas strong adhesion (ε = 5) generates cohesive but partially mislocalized aggregates. (B) Steady-state localization η_ss_(mean ± SEM, n = 8). (C) Mean tracking error ε_track_(t). (D) Representative snapshots at t = 300 s. Moderate cohesion stabilizes localization near the source, while excessive cohesion promotes off-target clustering. Parameters: τ_c_ = 5 s, A_noise_ = 0.35, N = 200, L = 600 μm.

### 3.3 Phase diagram of collective localization under temporally correlated noise

To quantify how isotropic cohesion modulates chemotactic performance under fluctuating environments, we systematically explored the two-dimensional parameter space defined by the noise correlation time *τ*_*c*_and the adhesion strength *ε*. For each condition, multiple independent realizations were simulated and the steady-state localization efficiency *η*_*ss*_ was computed. The resulting phase diagram (Fig. 2) reveals that collective localization depends strongly on interaction strength and weaker on noise correlation time within the explored regime. In the absence of cohesion (*ε* = 0), localization remains moderate but progressively deteriorates as temporal correlations increase, reflecting the reduced reliability of temporal gradient sensing under persistent fluctuations. Introducing isotropic adhesion systematically reshapes this behavior. Weak-to-intermediate cohesion (*ε* = 0.5–2) enhances localization across nearly the entire *τ*_*c*_range by suppressing noise-driven dispersion and stabilizing bacterial residence near the signal source. The strongest enhancement occurs at intermediate adhesion strength (*ε* = 2), where localization reaches *η*_*ss*_ ≈ 0.6–0.7over a broad range of correlation times. In this regime, cohesion robustly stabilizes collective occupation of the target while preserving sufficient mobility for chemotactic adaptation. Importantly, the collective rescue is strongly non-monotonic. At high adhesion (*ε* = 5), localization decreases despite the formation of highly cohesive aggregates. Representative snapshots show that strongly adhesive populations frequently collapse into compact off-target clusters that remain mechanically stable but only weakly coupled to the chemotactic landscape. This behavior reveals a partial decoupling between structural cohesion and functional localization: increasing connectivity does not necessarily improve targeting performance. The localization curves *η*_*ss*_(*τ*_*c*_)further illustrate this competition between collective stabilization and dynamical trapping. Intermediate cohesion extends the range over which localization remains robust under temporally correlated noise, whereas excessive adhesion reduces responsiveness by mechanically stabilizing mislocalized clusters. The system therefore exhibits an optimal interaction window balancing noise suppression and collective adaptability. Taken together, these results demonstrate that isotropic cohesion reshapes the limits of chemotactic performance under temporally structured fluctuations through competing effects of noise persistence, interaction-driven stabilization, and spatial miss-localization. The resulting collective behavior is governed not simply by the presence of cohesion, but by how interaction strength modulates the balance between structural stability and functional responsiveness.

**Figure 2.**
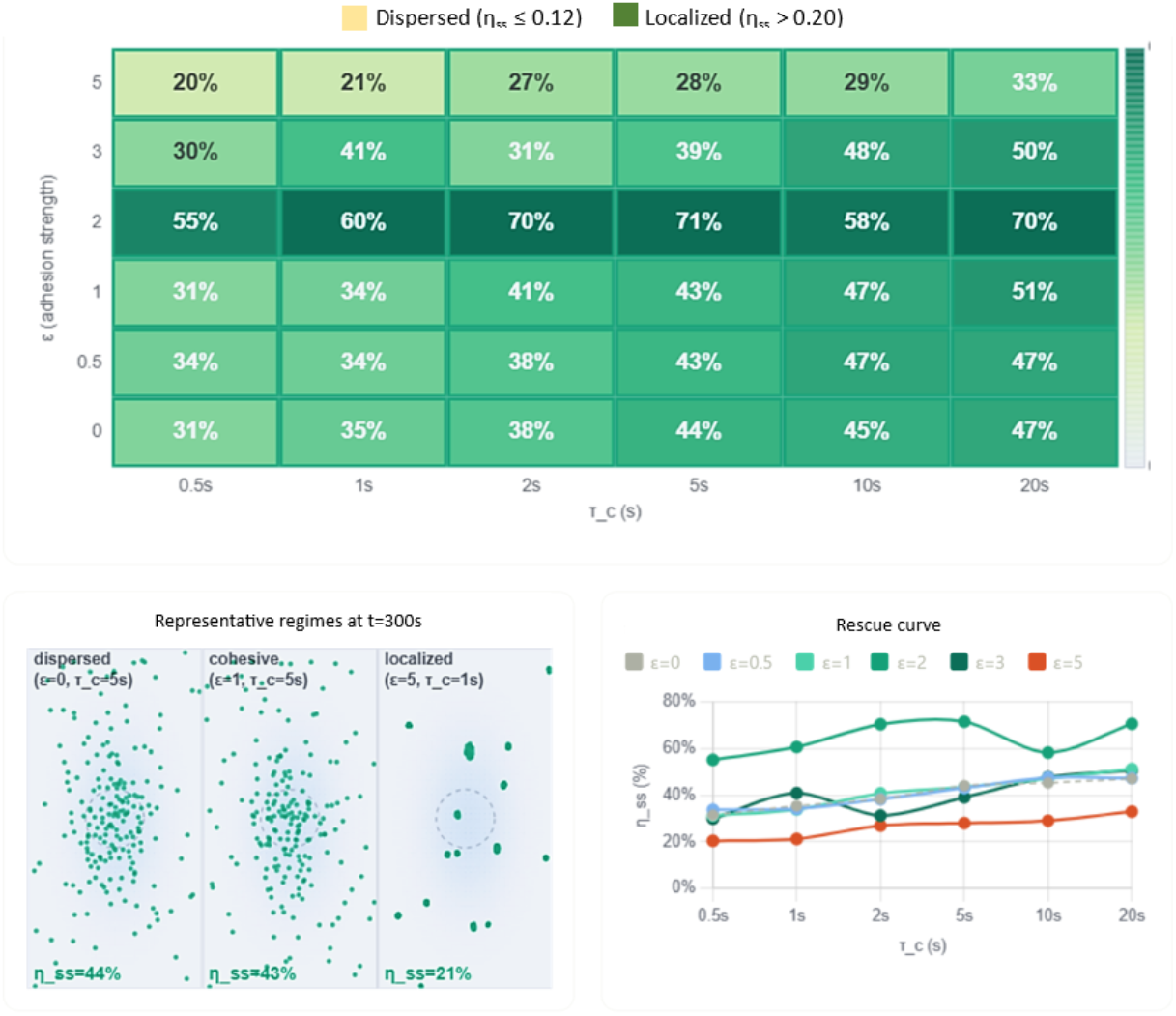
Phase diagram of cohesion-mediated chemotactic rescue. Top: Heatmap of steady-state localization efficiency η_ss_as a function of noise correlation time τ_c_and adhesion strength ε. Intermediate cohesion (ε = 2) maximizes localization robustness across a broad range of temporal noise conditions, whereas excessive cohesion (ε = 5) reduces performance through off-target clustering. Bottom left: Representative snapshots at t = 300 sshowing weakly cohesive, cohesive, and strongly adhesive regimes. Bottom right: Localization curves η_ss_(τ_c_)for different adhesion strengths. Cohesion enhances localization non-monotonically, revealing an optimal interaction window for collective chemotactic stabilization.

### 3.4 Selective interactions reorganize collective chemotaxis without enhancing localization

To investigate how interaction topology affects collective organization, we extended the model to a binary red– blue population with selective interactions. In this framework, heterotypic (R–B) pairs experience attractive forces, whereas homotypic (R–R and B–B) interactions are purely repulsive. This interaction hierarchy promotes cross-type bonding while suppressing same-type aggregation. We first compared the macroscopic chemotactic dynamics of the selective system with those of the isotropic adhesion model under identical environmental conditions. As shown in Fig. 3A, the localization dynamics *η*(*t*)are nearly indistinguishable between the two interaction architectures. Both systems reach comparable steady-state localization levels (*η*_*ss*_ ≈ 0.4–0.45), indicating that introducing interaction selectivity does not substantially modify global chemotactic performance.

**Figure 3.**
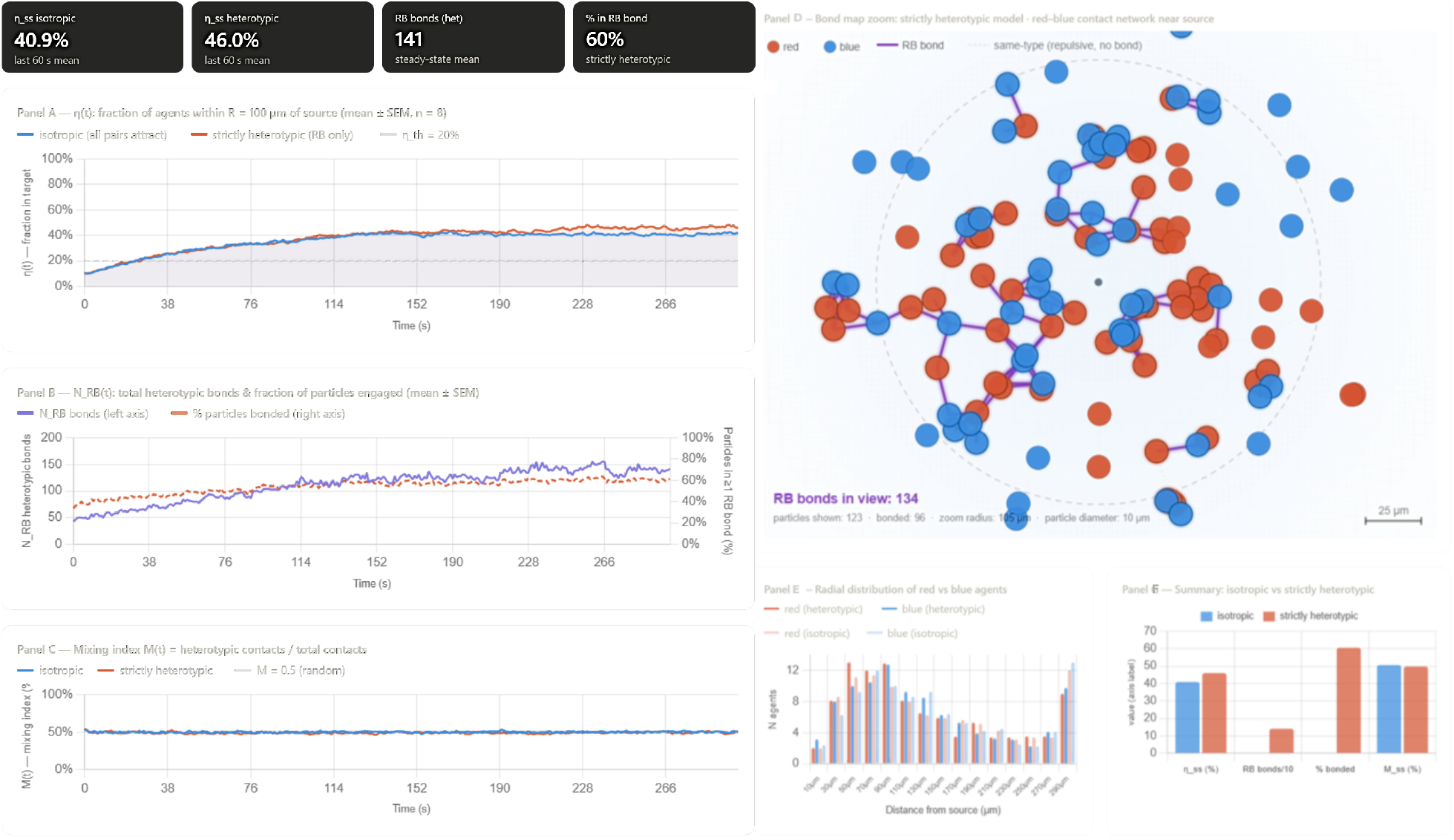
Selective interactions preserve localization while restructuring internal organization. (A) Localization dynamics η(t)for isotropic and selective interaction models under identical noise conditions. Both systems exhibit comparable steady-state localization. (B) Time evolution of heterotypic bond number N_RB_and fraction of bonded particles. (C) Mixing index M(t), remaining consistently above the random baseline (M = 0.5). (D) Representative snapshots and bond maps showing preferential red–blue organization within the localized population. Purple lines indicate active heterotypic bonds. (E) Distribution of heterotypic bond distances. (F) Summary comparison of bond statistics and mixing metrics. Selective interactions preserve collective localization while strongly reorganizing local contact topology.

Despite this similarity at the functional level, the internal structure of the population changes qualitatively. In the isotropic model, adhesion generates compact cohesive clusters with no internal compositional organization beyond spatial proximity. By contrast, selective interactions reorganize the collective into distributed heterotypic contact networks in which red and blue bacteria become preferentially interspersed throughout the localized region (Fig. 3D). To quantify this structural reorganization, we analyzed the statistics of heterotypic bonds. A bond was defined whenever a red–blue pair remained within the interaction radius *r*_*bond*_. The number of heterotypic contacts increased steadily during aggregation and saturated at approximately *N*_*RB*_ ≈ 140bonds at steady state (Fig. 3B). Simultaneously, nearly 60% of particles participated in at least one heterotypic bond, demonstrating that selective interactions generate an extended mixed-contact network spanning a large fraction of the population.

The mixing index *M*remained consistently above the random baseline throughout the simulation (Fig. 3C), confirming a persistent bias toward heterotypic organization. Importantly, this enhanced mixing emerged without significant changes in localization efficiency. The selective system therefore preserves global chemotactic function while substantially reorganizing local contact topology.

Representative snapshots and bond maps further illustrate this distinction (Fig. 3D–F). Rather than forming homogeneous condensates, the selective population develops dynamically maintained networks of transient red–blue contacts distributed throughout the cluster core. Same-type repulsion continuously frustrates unrestricted condensation, preventing complete structural collapse while maintaining strong local heterotypic connectivity. Together, these results establish that interaction topology acts as an independent organizational control parameter in collective chemotaxis. While isotropic cohesion primarily stabilizes localization through global condensation, selective interactions restructure the internal architecture of the collective without substantially modifying macroscopic targeting performance. Collective function and collective structure therefore emerge as partially decoupled properties controlled by distinct aspects of the interaction network.

### 3.5 Selective interactions generate noise-robust transient heterotypic contacts

To determine whether selective heterotypic interactions remain stable under fluctuating environments, we systematically explored the dependence of the selective model on noise correlation time *τ*_*c*_and noise amplitude *A*_*noise*_at fixed interaction strength *ε*_*RB*_ = 2. Across the full parameter range explored, the system remained in a weak-binding regime dominated by transient red–blue contacts rather than macroscopic mixed assemblies (Fig. 4A). The number of heterotypic bonds remained approximately constant (*N*_*RB*_ ≈ 20–25) across nearly two orders of magnitude in *τ*_*c*_and over all tested noise amplitudes (Fig. 4B, D). Similarly, the fraction of particles participating in at least one heterotypic bond remained stable (*f*_*RB*_ ≈ 0.24–0.28), indicating that local contact formation is largely insensitive to environmental fluctuation statistics. The mixing index also remained consistently close to, but slightly above, the random-contact baseline (*M* ≈ 0.50–0.54; Fig. 4C), demonstrating a persistent but modest bias toward heterotypic organization. Importantly, neither increasing noise persistence nor increasing fluctuation amplitude substantially altered the global structural state of the population. Representative snapshots further illustrate this behavior (Fig. 4E–G). Under low, intermediate, and high noise conditions, the population consistently formed dispersed red–blue dimers and small mixed-contact motifs distributed throughout the domain. Although heterotypic bonds remained dynamically stable, the system did not undergo large-scale condensation or percolating mixed-cluster formation. Interestingly, the number of heterotypic bonds formed near the signal source remained slightly elevated at low noise amplitudes (Fig. 4D), suggesting that selective contacts are preferentially stabilized in regions where chemotactic guidance remains partially effective. However, this effect remained local and did not propagate into macroscopic collective organization. Together, these results identify *ε*_*RB*_ = 2as a noise-robust transient-contact regime in which selective interactions continuously reorganize local contact topology without producing cohesive mixed assemblies. Unlike isotropic cohesion, which promotes collective condensation, weak selective interactions primarily bias short-range contact statistics while preserving globally dispersed population structure.

**Figure 4.**
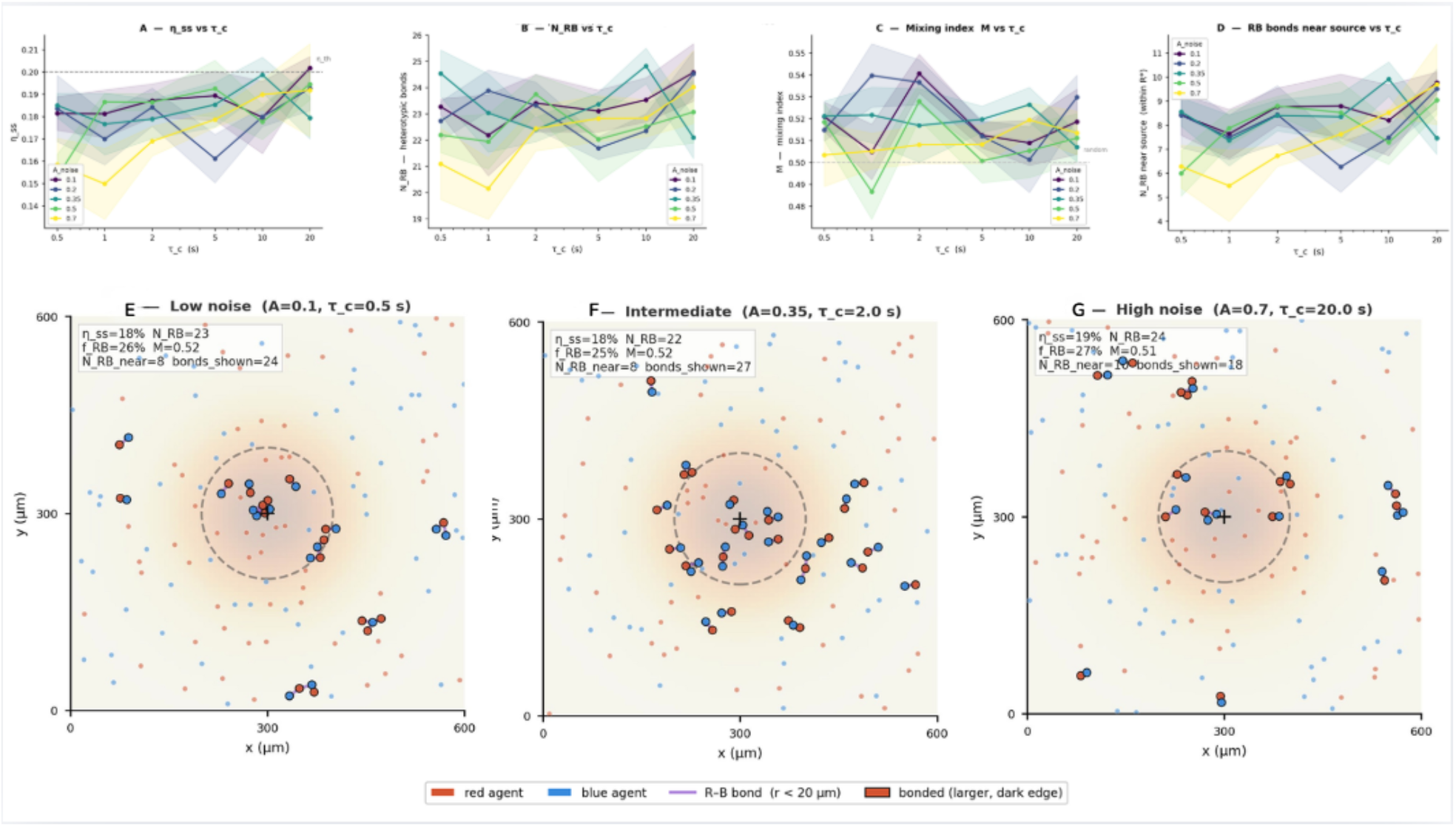
Noise robustness of transient heterotypic contacts in the selective red–blue model. Steady-state statistics of the selective interaction model under variations in noise correlation time τ_c_and noise amplitude A_noise_. (A) Steady-state localization η_ss_as a function of τ_c_. (B) Number of heterotypic red–blue bonds N_RB_. (C) Mixing index M; dashed line indicates the random-contact baseline M = 0.5. (D) Number of heterotypic bonds formed within the target region. (E–G) Representative snapshots at t = 120 sfor low, intermediate, and high noise conditions. Purple lines indicate active R–B bonds (r_ij_ < 20 μm); bonded agents are shown with larger disks and dark edges. Selective interactions generate persistent transient heterotypic contacts across all noise conditions but do not produce macroscopic mixed condensation. Parameters: S_0_ = 800, ε_RB_ = 2, a = 0.10 μm^−1^, r_0_ = 16 μm, N = 150, L = 600 μm, T = 120 s, n_rep_ = 4.

### 3.6 Phase diagram of selective contact organization

To determine whether selective heterotypic interactions generate robust collective assemblies under temporally correlated noise, we systematically explored the parameter space defined by the heterotypic interaction strength *ε*_*RB*_and the environmental correlation time *τ*_*c*_. Unlike the isotropic cohesion model, where increasing interaction strength promoted macroscopic clustering and collective localization, the selective model exhibited a qualitatively distinct phenomenology dominated by distributed heterotypic contact networks rather than large-scale condensation. Across the explored parameter range, the largest-cluster fraction remained low (*C*_*max*_ ≈ 0.13–0.16), indicating that selective heterotypic attraction alone is insufficient to generate cohesive mixed assemblies under the present interaction architecture. Instead, the population remained in a structurally dispersed regime composed primarily of transient dimers, trimers, and small mixed-contact motifs distributed throughout the domain.

In contrast, local structural observables varied systematically with interaction strength. Increasing *ε*_*RB*_progressively enhanced the number of heterotypic bonds *N*_*RB*_, which increased from approximately 26 at *ε*_*RB*_ = 0to more than 30 at *ε*_*RB*_ = 30. Similarly, the mixing index increased from values near the random baseline (*M* ≈ 0.50) to *M* ≈ 0.56, demonstrating a robust bias toward heterotypic organization. Importantly, these changes occurred without a corresponding increase in macroscopic cohesion, revealing a decoupling between contact selectivity and large-scale aggregation.

A notable result is the weak dependence of these structural observables on the temporal correlation time *τ*_*c*_. Although temporally correlated noise strongly affected localization in the isotropic model, the selective system maintained similar values of *N*_*RB*_, *M*, and *C*_*max*_across a broad range of noise persistence times. This indicates that local interaction topology dominates over environmental fluctuations in determining collective structure. Representative snapshots illustrate this progression (Fig. 5). In the absence of selective attraction (*ε*_*RB*_ = 0), particles remain weakly correlated apart from chemotactic accumulation near the source. Increasing *ε*_*RB*_produces progressively denser networks of transient red–blue bonds, visible as distributed heterotypic contact motifs throughout the domain. However, even at the highest interaction strengths explored, these contacts do not collapse into compact mixed condensates. Instead, same-type repulsion continuously frustrates large-scale aggregation, stabilizing a dynamically maintained contact-network state.

**Figure 5.**
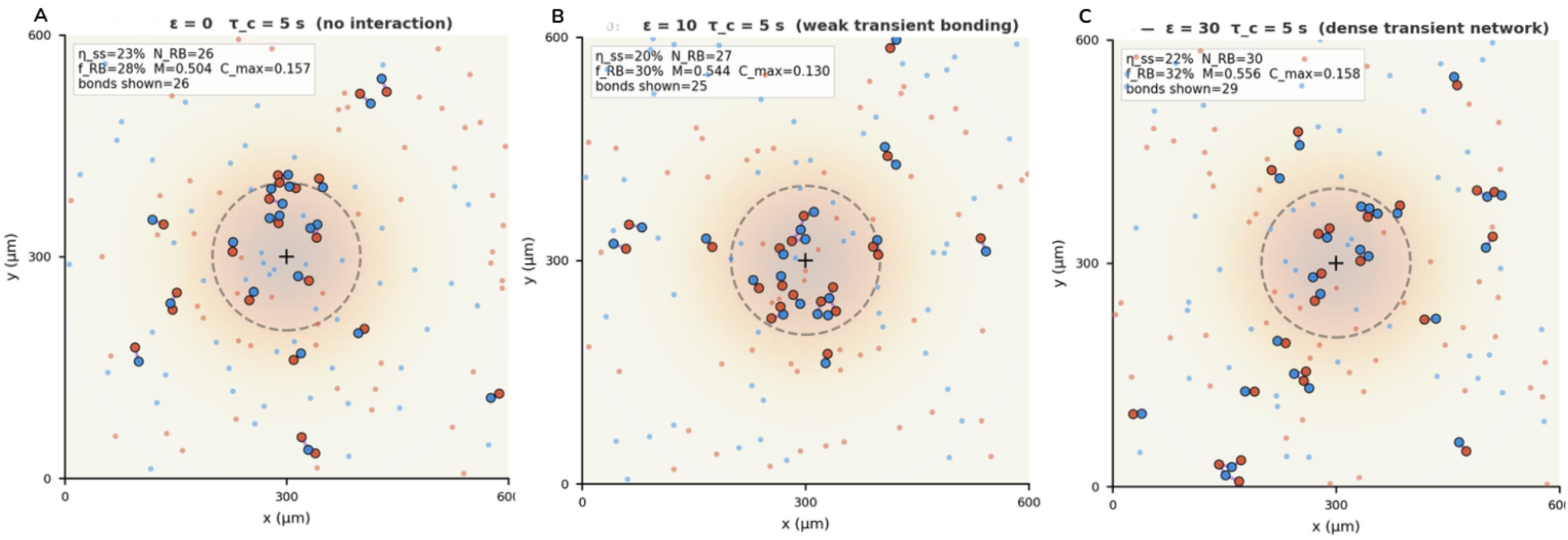
Selective interaction strength reorganizes contact topology without inducing macroscopic condensation. Representative steady-state configurations of the selective red–blue model for increasing heterotypic interaction strength *ε*_*RB*_at fixed *τ*_*c*_ = 5 s. Red–blue pairs within the bonding distance are connected by purple lines; bonded agents are shown with larger disks and dark edges. Dashed circles indicate the localization region around the signal source. (A) *ε*_*RB*_ = 0: weakly correlated population with near-random mixing. (B) *ε*_*RB*_ = 10: emergence of transient heterotypic dimers and mixed-contact motifs. (C) *ε*_*RB*_ = 30: dense transient heterotypic contact network without macroscopic condensation.

Together, these results demonstrate that selective interaction architectures reorganize the topology of inter-agent contacts without inducing macroscopic collective condensation. Whereas isotropic adhesion promotes cohesive localization through global aggregation, selective heterotypic interactions generate structurally dispersed but topologically organized mixed-contact networks. The emergent collective state is therefore not a condensed assembly phase, but a frustrated active-matter regime whose organization is governed primarily by interaction specificity rather than by environmental noise statistics.

### 3.7 Comparison between isotropic and selective systems

To isolate the role of interaction topology, isotropic and selective systems were compared under identical chemotactic and environmental noise conditions. Both models shared the same run-and-tumble dynamics, Ornstein–Uhlenbeck noise statistics, signal landscape, and comparable interaction scales, differing only in the structure of the pairwise interaction network. Despite their fundamentally different interaction architectures, both systems exhibited similar levels of steady-state localization (Fig. 6A). Across the explored interaction strengths, *η*_*ss*_remained within overlapping ranges for isotropic and selective models, indicating that interaction topology does not strongly modify the overall ability of the population to accumulate near the signal source.

**Figure 6.**
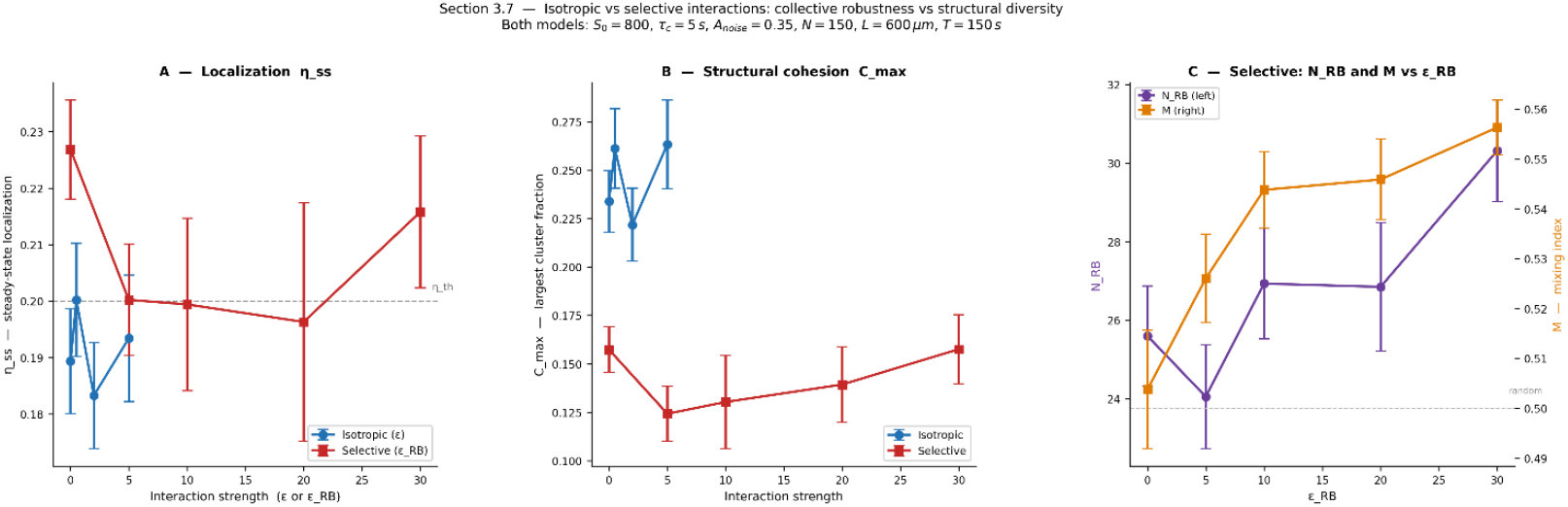
Interaction topology decouples collective function from collective architecture. Comparison between isotropic and selective interaction models under identical chemotactic and environmental noise conditions (τ_c_ = 5 s, A_noise_ = 0.35). (A) Steady-state localization η_ss_as a function of interaction strength. Both models exhibit comparable localization levels despite distinct interaction architectures. Dashed line indicates the localization threshold η_th_. (B) Largest-cluster fraction C_max_. Isotropic interactions generate substantially larger cohesive assemblies than selective interactions. (C) Selective-model structural observables as a function of ε_RB_: heterotypic bond number N_RB_(purple) and mixing index M(orange). Increasing selective attraction enhances heterotypic organization without inducing macroscopic condensation. Together, the results demonstrate that interaction topology strongly modifies collective structure while only weakly affecting global localization performance.

In contrast, structural observables differed markedly between the two interaction schemes. The isotropic model consistently produced larger connected clusters, with *C*_*max*_ ≈ 0.22–0.26, reflecting the emergence of cohesive condensate-like assemblies (Fig. 6B). The selective model, however, remained in a structurally dispersed regime with *C*_*max*_ ≈ 0.13–0.16even at high *ε*_*RB*_, demonstrating that heterotypic attraction combined with homotypic repulsion prevents macroscopic condensation.

At the same time, selective interactions strongly reorganized local contact topology. Increasing *ε*_*RB*_progressively enhanced the number of heterotypic bonds *N*_*RB*_and increased the mixing index from values near the random baseline (*M* ≈ 0.50) to *M* ≈ 0.56(Fig. 6C). Thus, while isotropic cohesion primarily reinforces global connectivity, selective interactions instead promote distributed mixed-contact organization. These results demonstrate that collective function and collective structure constitute partially independent properties of the system. Similar localization efficiencies can emerge from fundamentally different organizational states: compact cohesive condensates in the isotropic model, or spatially dispersed heterotypic contact networks in the selective model. Interaction topology therefore acts as an independent control parameter governing how collective organization is physically realized.

Together, this comparison establishes a central result of LUCA2: collective robustness and collective architecture are not uniquely coupled. Populations with comparable chemotactic performance may nevertheless exhibit profoundly different internal organization depending on the topology of inter-agent interactions.

## 4. Results

### 4.1 Collective rescue as a non-equilibrium phase transition

The results presented here demonstrate that collective chemotactic localization under temporally correlated noise cannot be understood as a simple extension of single-cell sensing. When the environmental correlation time exceeds the characteristic run time (*τ*_*c*_ > *τ*_*run*_), temporal comparisons become unreliable and isolated bacteria fail to maintain stable localization despite the presence of a persistent mean gradient. Under these conditions, chemotactic failure emerges not from the absence of directional information, but from the persistence of fluctuations over timescales longer than the cellular sensing cycle. A central result of this work is that short-range isotropic cohesion fundamentally alters this limitation. Interactions do not directly improve gradient sensing at the single-agent level; instead, they generate a collective stabilization mechanism that suppresses noise-driven dispersion at the population scale. Localization therefore emerges as an emergent property of the interacting ensemble rather than as the sum of independent chemotactic trajectories.

This transition can be interpreted as a form of non-equilibrium collective rescue. In the non-interacting regime, temporally correlated fluctuations continuously decorrelate the population from the target, preventing persistent accumulation near the source. Introducing cohesion changes the dynamical balance between stochastic dispersion and spatial retention. Bacteria that transiently approach the signal source form cohesive clusters stabilized by short-range attraction, increasing their effective residence time near the target region. In turn, these retained bacteria act as local nucleation centers that capture additional agents, amplifying accumulation through positive feedback.

Importantly, this mechanism operates without alignment interactions, explicit communication, or direct gradient forces. The collective does not emerge from coordinated decision-making, but from the interaction-driven filtering of stochastic fluctuations. Cohesion effectively performs a distributed averaging process: while individual trajectories remain noisy and unreliable, the interacting population suppresses dispersive fluctuations and stabilizes coherent occupation of the target region. In this sense, the collective behaves as a dynamical noise filter whose robustness exceeds that of its constituent agents. This behavior closely resembles collective averaging phenomena in active matter systems, where inter-agent interactions suppress individual fluctuations and generate emergent directed transport. However, unlike canonical flocking or alignment-based models, the present system achieves collective stabilization purely through isotropic cohesion combined with chemotactic bias. The resulting localized phase therefore emerges without orientational order or explicit collective steering. The transition also shares conceptual similarities with non-equilibrium phase transitions in interacting active particles. Above a finite interaction threshold, the system transitions from a dispersed phase to a collectively stabilized localized phase in which fluctuations become dynamically confined. However, the present system differs fundamentally from conventional motility-induced phase separation because localization is externally biased by a chemotactic landscape and functionally defined through target occupancy rather than density alone. The relevant transition is therefore not simply structural condensation, but the emergence of a functionally localized phase.

Our results further show that cohesion and localization are not equivalent observables. Strong interactions can stabilize cohesive clusters that remain spatially displaced from the source, generating structurally ordered yet functionally ineffective states. The existence of this cohesive but non-localized regime demonstrates that structural stabilization and functional localization constitute distinct collective transitions controlled by separate mechanisms. Collective robustness emerges only when connectivity and environmental bias become dynamically coupled. Taken together, these findings suggest that collective chemotactic behavior under temporally structured noise should be understood as an emergent non-equilibrium phenomenon in which interactions reshape the effective information-processing capacity of the population. Rather than merely stabilizing spatial organization, cohesion creates a new collective dynamical regime in which localization becomes possible beyond the limits of individual sensing.

### 4.2 Decoupling between cohesion and function

A key conceptual outcome of this study is that collective cohesion and functional chemotactic localization are not equivalent phenomena. Although attractive interactions can stabilize spatially coherent clusters, the existence of a cohesive aggregate does not necessarily imply successful targeting of the signal source. Our simulations reveal that structural order and functional order constitute distinct dynamical observables that can become partially or fully decoupled depending on the interaction regime. This distinction becomes particularly evident at high adhesion strengths. Moderate isotropic cohesion enhances localization by increasing the residence time of bacteria near the target region and suppressing noise-induced dispersion. However, when cohesion becomes too strong, the collective increasingly behaves as a mechanically self-stabilized object whose dynamics are dominated by internal connectivity rather than chemotactic responsiveness. In this regime, clusters remain cohesive even when displaced away from the source, producing structurally ordered yet functionally ineffective states. The population therefore transitions from a chemotactically responsive collective to a mechanically trapped condensate.

The existence of these off-target cohesive states demonstrates that aggregation alone is insufficient to guarantee collective function. A cluster may exhibit high spatial coherence, low fragmentation, and persistent inter-agent connectivity while simultaneously failing to maximize occupancy of the target region. Structural condensation and chemotactic performance therefore emerge from partially competing dynamical tendencies: cohesion suppresses stochastic dispersion, but excessive cohesion also reduces the ability of the population to reorganize in response to environmental information. From a non-equilibrium perspective, this behavior resembles dynamical arrest phenomena observed in active condensed matter systems. Increasing interaction strength progressively shifts the dominant timescale of the system from chemotactic reorientation toward collective mechanical stabilization. Beyond an intermediate regime, the collective loses adaptive flexibility and becomes increasingly insensitive to the gradient landscape. Functional optimization therefore occurs not at maximal cohesion, but near an intermediate interaction regime where structural stability and environmental responsiveness remain balanced.

Importantly, this decoupling provides a mechanistic explanation for the phase structure observed in the isotropic model. The dispersed phase corresponds to insufficient connectivity, where fluctuations dominate and localization fails. The localized phase emerges when cohesion is strong enough to stabilize occupancy near the source while preserving collective mobility. At still larger interaction strengths, however, structurally cohesive but functionally suboptimal states become increasingly probable. The system therefore exhibits a sequential organization hierarchy: dispersion, functional localization, and finally mechanically dominated cohesion. This finding has direct implications for the interpretation of cohesion in biological systems. Cellular adhesion is often assumed to enhance collective navigation simply by maintaining group integrity. The present results demonstrate that this assumption is not generically correct: cohesion enhances function only within a finite window of interaction strengths, above which structural order and functional order become decoupled. In this sense, collective robustness does not emerge from aggregation alone, but from the balance between connectivity and environmental responsiveness. More generally, the results suggest that collective chemotaxis should not be characterized solely through geometric observables such as cluster size or density. Functional observables— including target occupancy, localization stability, and responsiveness to fluctuating environments—capture independent aspects of the collective state. From a physics perspective, this decoupling is analogous to frustration phenomena in condensed matter systems, where strong local order does not necessarily imply global functional order. Similarly, strong short-range cohesion here does not guarantee successful collective localization. Taken together, these findings establish that collective structure and collective function constitute partially independent organizational principles in active chemotactic matter. The emergence of functionally localized states therefore depends not only on whether agents aggregate, but on how aggregation dynamically couples to environmental information under non-equilibrium conditions.

### 4.3 Interaction topology controls collective architecture

The comparison between isotropic and selective interaction models demonstrates that collective organization is governed not only by interaction strength, but also by interaction topology. Although both models operate under identical chemotactic dynamics, environmental noise statistics, and comparable interaction scales, they generate qualitatively different collective architectures. Isotropic interactions drive macroscopic condensation and cohesive localization, whereas selective interactions reorganize local contact structure while actively preventing large-scale aggregation.

In the isotropic model, every pairwise interaction contributes cooperatively to global cohesion. Because attraction is independent of agent identity, local contacts reinforce one another and promote the formation of compact collective condensates once the interaction strength exceeds a threshold. The resulting structures are mechanically cohesive and spatially compact, enabling collective stabilization against temporally correlated fluctuations. Condensation therefore emerges naturally from the positive feedback between aggregation and retention near the signal source.

The selective model operates according to fundamentally different principles. Heterotypic attraction promotes red–blue contact formation, while homotypic repulsion continuously destabilizes same-type clustering. These competing interaction rules impose a topological constraint on assembly: for a large mixed aggregate to form, red and blue agents must remain interleaved across multiple spatial scales. Same-type repulsion prevents the unrestricted growth of cohesive domains and continuously fragments incipient aggregates, frustrating macroscopic condensation.

As a consequence, the system does not collapse into dense condensates but instead stabilizes distributed mixed-contact networks composed primarily of transient dimers, trimers, and short-lived heterotypic motifs dispersed throughout the domain. Importantly, these structures remain statistically robust across broad ranges of environmental noise despite the absence of global assembly. The weak dependence of mixing observables on the noise correlation time indicates that local contact topology is governed primarily by interaction architecture rather than by chemotactic performance itself.

This distinction highlights a broader principle in active matter systems: interaction topology can independently control collective architecture even when macroscopic function remains largely unchanged. In isotropic systems, connectivity reinforces condensation and produces globally coherent phases. In selective systems, connectivity becomes geometrically constrained, generating frustrated active-matter states in which local ordering persists without large-scale structural collapse. The collective therefore remains dynamically organized, but the organization is encoded in the topology of contacts rather than in bulk density accumulation. From a physical perspective, this frustrated regime resembles competing-interaction systems studied in soft condensed matter and multicomponent assembly, where short-range attraction combined with longer-range or selective repulsion generates arrested phases, gels, or microstructured states instead of complete phase separation. The present results suggest that analogous frustration mechanisms can emerge in active chemotactic systems, stabilizing transient contact-network states with well-defined statistical organization.

The selective model therefore represents a form of programmable active organization. By modifying only the interaction matrix—without altering motility, sensing, or environmental forcing the system transitions from cohesive condensates to distribute mixed-contact networks. This demonstrates that collective architecture can be tuned independently of the underlying chemotactic dynamics, suggesting that local interaction rules constitute a powerful control layer for engineering active biological matter.

More broadly, these findings suggest that collective biological organization may often arise through topological regulation of local interactions rather than through global condensation alone. Systems with selective adhesion, receptor specificity, or identity-dependent binding may exploit frustration to maintain structural diversity, distributed connectivity, and adaptive reconfigurability simultaneously. In this framework, isotropic cohesion and selective interactions do not represent stronger or weaker versions of the same collective mechanism, but rather two distinct organizational paradigms with fundamentally different physical consequences.

### 4.4 Biological and synthetic implications

The mechanisms identified in this work have implications that extend beyond bacterial chemotaxis itself, touching broader questions in collective biological organization, synthetic active matter, and programmable living systems. Our results show that relatively simple interaction rules can qualitatively reshape the robustness, structure, and functionality of populations navigating fluctuating environments. In this sense, collective behavior emerges not only from sensing capabilities, but from the interaction architecture through which populations redistribute information and stabilize spatial organization. In biological systems, temporally fluctuating chemical landscapes are the rule rather than the exception. Microbial communities, immune cells, and migrating multicellular populations routinely operate in heterogeneous environments where signals are transient, spatially fragmented, and dynamically perturbed by diffusion, transport processes, metabolic consumption, or competing consumers. Under such conditions, the ability of populations to maintain coherent localization despite unreliable local information may critically depend on collective stabilization mechanisms analogous to those identified here.

For bacterial consortia and biofilm-forming systems, the isotropic cohesion regime provides a minimal physical mechanism through which surface adhesion, extracellular polymers, or matrix-mediated interactions can enhance robustness against environmental uncertainty. Even weak attractive interactions can transform populations from dispersed exploratory states into collectively stabilized localized phases. Importantly, this stabilization emerges without explicit communication, quorum sensing, or centralized coordination, indicating that purely mechanical or adhesive coupling may substantially extend the functional limits of chemotactic populations under temporally structured noise. The selective interaction model points toward a distinct organizational principle relevant to multispecies microbial communities and heterogeneous multicellular assemblies. In natural consortia, inter-species adhesion is often mediated by complementary surface lectins, lipopolysaccharides, extracellular peptides, or receptor–ligand specificity—precisely the type of selective heterotypic interaction represented in the model. Our results predict that such interactions do not necessarily enhance global chemotactic targeting, but instead reorganize the internal topology of contacts, generating distributed mixed-contact networks whose structure remains robust across environmental fluctuations.

Such architecture may be advantageous in systems where local compositional diversity is functionally important. Cross-feeding microbial consortia, cooperative metabolic networks, and spatially distributed signaling communities frequently require sustained proximity between distinct cell types while simultaneously avoiding complete macroscopic condensation. The emergence of transient heterotypic motifs without large-scale aggregation therefore suggests a plausible physical mechanism through which heterogeneous biological systems maintain both connectivity and spatial flexibility in fluctuating environments. These ideas also connect naturally to the design of synthetic living materials and programmable active matter. Modern synthetic biology increasingly enables cells to be engineered with tunable adhesion molecules, synthetic peptide pairs, receptor specificity, and dynamic interaction rules. Within this context, the present results suggest that modifying interaction topology alone may suffice to switch between qualitatively distinct collective states: cohesive condensates, distributed contact networks, or dynamically frustrated assemblies. Interaction architecture therefore becomes a programmable design variable for controlling emergent organization at the population scale.

The decoupling identified here between structural cohesion and functional localization provides a particularly relevant design principle. By tuning interaction strength, one can independently regulate whether a population forms macroscopic aggregates (high *C*_*max*_) or achieves robust localization near a target (high *η*_*ss*_). Likewise, selective interactions allow internal contact organization to be programmed without substantially modifying macroscopic chemotactic performance. Structural diversity and collective robustness therefore emerge as partially independent targets. The phase diagrams further establish the environmental correlation time *τ*_*c*_as a key control parameter for collective robustness. In engineered microfluidic environments or synthetic ecological platforms, where signal dynamics can be externally programmed, temporal noise structure becomes an experimentally tunable variable rather than uncontrolled perturbation. Environments with *τ*_*c*_ < *τ*_*run*_permit robust individual chemotaxis because rapid fluctuations are efficiently averaged out, whereas environments with *τ*_*c*_ ≫ *τ*_*run*_require collective interactions to sustain localization. This provides a physically grounded framework for controlling microbial navigation in diagnostics, targeted delivery, and spatially structured biosystems.

The implications also extend beyond biological matter. In swarm robotics and distributed autonomous systems, individual agents frequently operate under noisy or incomplete environmental information. The collective rescue mechanism identified here suggests that local coupling rules can compensate for unreliable sensing at the agent level, enabling robust collective targeting even when individual guidance fails. Conversely, selective interaction rules may provide a route toward maintaining distributed yet coordinated swarms without global clustering, which may be advantageous for exploration, adaptive coverage, or decentralized transport.

More broadly, our results support a view of collective organization as a form of emergent computation performed through interaction dynamics. Cohesion, selectivity, and noise filtering collectively reshape the effective information-processing capacity of the population, determining whether environmental information becomes amplified, suppressed, or spatially redistributed. From this perspective, active populations are not simply collections of motile particles, but non-equilibrium systems whose interaction architecture defines the class of collective functionality they can realize. Finally, the distinction identified here between functional localization and structural organization may prove broadly relevant across active and living matter systems. Biological collectives rarely optimize a single objective; robustness, adaptability, diversity, and spatial coherence often compete depending on environmental context. The coexistence of dispersed, localized, cohesive, and frustrated organizational regimes suggests that many collective biological phenomena may arise from navigating these competing constraints rather than from maximizing aggregation alone.

## 5. Conclusions

In this work, we presented a two-dimensional agent-based model of interacting with run-and-tumble bacteria navigating temporally correlated chemical environments. By combining temporal chemotactic sensing with tunable inter-agent interactions, the model establishes a unified framework for studying how environmental memory and interaction topology jointly shape collective behavior under noise. Our results demonstrate first that temporally correlated fluctuations impose a fundamental limit on individual chemotaxis. When the environmental correlation time exceeds the intrinsic run timescale (*τ*_*c*_ > *τ*_*run*_), temporal comparisons become unreliable and non-interacting populations fail to sustain stable localization despite the presence of a persistent mean gradient. Chemotactic failure therefore emerges primarily from the temporal persistence of fluctuations rather than from noise amplitude alone. Second, we show that isotropic short-range cohesion partially rescues collective chemotactic function by suppressing noise-driven dispersal and stabilizing occupancy near the signal source. Importantly, this rescue is non-monotonic. Increasing adhesion strength enhances structural connectivity (*C*_*max*_) more robustly than localization efficiency (*η*_*ss*_), revealing that cohesion and function are distinct collective observables with separate onset conditions. The emergence of cohesive but weakly localized states demonstrates that aggregation alone is insufficient to guarantee collective targeting.

Third, selective heterotypic interactions generate a qualitatively different organizational regime. Rather than producing macroscopic mixed condensates, heterotypic attraction combined with homotypic repulsion stabilizes transient red–blue contact networks composed of dimers and small mixed motifs. These structures remain robust across broad ranges of environmental noise conditions, indicating that local contact topology is governed primarily by interaction specificity rather than by global chemotactic performance.

Taken together, these findings establish interaction topology as an independent control parameter for collective active matter organization. Isotropic and selective interactions produce comparable localization efficiencies while generating fundamentally different structural states: cohesive condensates in the isotropic case and frustrated mixed-contact networks in the selective case. Collective robustness and collective structural diversity therefore emerge as partially competing organizational principles controlled by the architecture of inter-agent interactions. More broadly, this research connects chemotaxis under fluctuating environments with concepts from non-equilibrium statistical physics and programmable active matter. The framework provides experimentally testable predictions for engineered bacterial consortia, synthetic living materials, and microfluidic systems in which temporal signal structure and interaction specificity can be independently tuned. More generally, the results suggest that collective biological function may emerge not solely from improved sensing, but from the dynamical regulation of interactions that redistribute and stabilize information at the population scale.

## Notes

### Competing Interest Statement

The authors have declared no competing interest.

### Funding

- European Union: EIC Pathfinder project THOR (Grant Agreement number 101099719), EIC Pathfinder project ISOS (Grant Agreement number 101130454)
- Comunidad de Madrid: B2017/BMD-3760 Neurocentro-CM, P022/BMD-7236 MINA-CM

